# Type IV CRISPR-Cas systems are highly diverse and involved in competition between plasmids

**DOI:** 10.1101/780106

**Authors:** Rafael Pinilla-Redondo, David Mayo-Muñoz, Jakob Russel, Roger A. Garrett, Lennart Randau, Søren J. Sørensen, Shiraz A. Shah

## Abstract

CRISPR-Cas systems provide prokaryotes with adaptive immune functions against viruses and other genetic parasites by leveraging small non-coding RNAs for nuclease-dependent degradation of their nucleic acid targets. In contrast to all other types of CRISPR-Cas systems, the mechanisms and biological roles of type IV systems have remained largely overlooked. Here, we describe a previously uncharted diversity of type IV gene cassettes, distributed across diverse prokaryotic genome backgrounds, and propose their classification into subtypes and variants. Congruent with recent findings, type IV modules were primarily found on plasmid-like elements. Remarkably, via a comprehensive analysis of their CRISPR spacer content, these systems were found to exhibit a strong bias towards the targeting of other plasmids. Our data indicate that the functions of type IV systems have diverged from those of other host-related CRISPR-Cas immune systems to adopt a yet unrecognised role in mediating conflicts between plasmids that compete to monopolize their hosts. Furthermore, we find evidence for cross-talk between certain type IV and type I CRISPR-Cas systems that co-exist intracellularly, thus providing an answer to the enigmatic absence of adaptation modules in these systems. Collectively, our results lead to the expansion and reclassification of type IV systems and provide novel insights into the biological function and evolution of these elusive systems.

## Introduction

Clustered regularly interspaced short palindromic repeats (CRISPR), together with their CRISPR-associated (Cas) genes, constitute a diverse family of nucleic acid-based adaptive immune systems that protect archaea and bacteria against invading mobile genetic elements (MGEs). These defence systems are classified by virtue of their modular composition and structure, into two major groups, Class 1 and Class 2, that are respectively subdivided into types I, III, IV and types II, V, VI^1^.

Over the last decade, our knowledge regarding CRISPR-Cas systems has expanded at an exceptional rate, mainly driven by a strong effort to harness their biotechnological potential ^2–4^. To date, the functions and mechanisms of action of all known CRISPR-Cas types have been characterized in detail, except for type IV for which the biological function(s) remain enigmatic. Importantly, type IV CRISPR-Cas modules have recently been reported to be primarily encoded by plasmids or, occasionally, by prophage genomes, evidencing the recurrent transfer of the CRISPR-Cas machinery to and from MGEs^5^. Furthermore, although type IV cas operons are frequently associated with CRISPR arrays, they lack certain hallmark components of other CRISPR-Cas systems, including the highly conserved adaptation module and an effector nuclease^1^. Consequently, these reduced systems have been proposed to exhibit altered CRISPR-Cas functions or to be functionally defective ^6^.

To date, type IV CRISPR-Cas loci are classified into two distinct subtypes, IV-A and IV-B, both of which share a common set of effector module proteins, including a highly diverged Cas7 (Csf2), Cas5 (Csf3), and a smaller version of Cas8 (Csf1)^1^. Moreover, subtype IV-A loci encode a DinG family helicase (Csf4), a type IV-specific Cas6-like protein (Csf5), and they typically co-locate with a CRISPR array. In contrast, subtype IV-B loci lack *dinG*, *csf5* and an associated CRISPR array but they encode a putative “small subunit” (Cas11) and they often neighbour a *cysH* gene^7,8^. A recent structural and biochemical analysis of a subtype IV-A CRISPR-Cas system demonstrated the essential role of the Cas6-like enzyme in both the maturation of crRNAs and in the subsequent formation of a Cascade-like crRNA-guided effector complex, composed of Csf1, Csf3, Csf5 and multiple copies of Csf2^9^. These data suggest that the subtype IV-A effector complexes, as in other CRISPR-Cas systems, survey the cellular environment searching for matching nucleic acid targets. However, the study concluded that the spacers of the associated CRISPR arrays yielded no clear spacer-protospacer matches^9^, but an earlier larger-scale analysis reported putative sequence matches to MGEs of which 72% were reported to be of viral origin^10^.

In summary, it is plausible that subtype IV-A systems perform a defensive role, although the apparent absence of an effector nuclease suggests that the mechanism of interference differs significantly from those of other CRISPR-Cas systems. Consistent with this view, alternative functions have been suggested for type IV systems, including their involvement in plasmid propagation mechanisms, and in the enhancement of recombination events with other nucleic acids^7,9^. In particular, the absence of CRISPR arrays linked to the minimal subtype IV-B system provides support for the effector module machinery participating in alternative cellular functions^7^. In the present study we have undertaken a comparative genomics approach to survey all publicly available bacterial and archaeal genomes for type IV CRISPR-Cas systems. The collected type IV systems were then subjected to an in-depth bioinformatic characterisation to obtain insights into their exceptional biology and evolution.

## Results

### Expanding the number of identified type IV CRISPR-Cas systems

In order to perform a comprehensive analysis of the diversity and distribution of type IV CRISPR-Cas systems, we first sought to expand the repertoire of currently identified loci. Although Csf1 has been proposed as a signature protein for type IV systems^1^, we found that it was unsuitable, owing to its high level of sequence divergence between subtypes/variants and because of its absence from some loci. Instead, the Cas7-like (Csf2) protein was found to be the most conserved protein, and it was used as an initial query for searches against all publicly available complete and draft genomes (obtained from https://ftp.ncbi.nih.gov). Out of 883 detected Csf2 proteins (Supplementary Fig. 1), 69 diverse representatives were selected for further analysis. The gene neighbourhoods of these representatives were explored systematically and annotated manually via PSI-BLAST^11^ searches, protein clustering^12^ and profile-profile alignments^12,13^. An aggregate protein similarity tree was then generated including all proteins from the curated type IV modules. Finally, their corresponding gene maps were compared to gauge the diversity of their genetic compositions (Supplementary Fig. 2).

### Type IV systems display a previously uncharted diversity of loci architectures

Our phylogenetic analysis outlines a hitherto unrecognized richness of type IV gene arrangements and reveals a complex evolutionary relationship between the different variants, pervaded by clear instances of horizontal gene transfer (Fig. 1). The identified type IV loci are distributed across five major phylogenetically discrete groups that show consistent differences in their genetic compositions (Supplementary Fig. 3). Notably, a set of archaeal type IV modules were found to cluster as a clear outgroup and during the preparation of this work were proposed as a new subtype: IV-C (S.A.S personal communication with K.S. Makarova). These distinctive loci share key organizational features with type III CRISPR-Cas systems, including the presence of a Cas10-like protein in place of Csf1, their common association with type I CRISPR-Cas systems, and the frequent absence of CRISPR arrays and adaptation modules (Supplementary Fig. 4). Importantly, exhaustive protein domain searches with the IV-C Cas10-like protein revealed the typical Zn finger domain found in the middle section of other Class 1 CRISPR-Cas large subunits (Cas8, Csf1 and Cas10 families^14,15^) and, similarly to type III Cas10 proteins, an N-terminal HD nuclease domain that is suggestive of DNA cleavage activity (Supplementary Table 1).

**Fig. 1.**
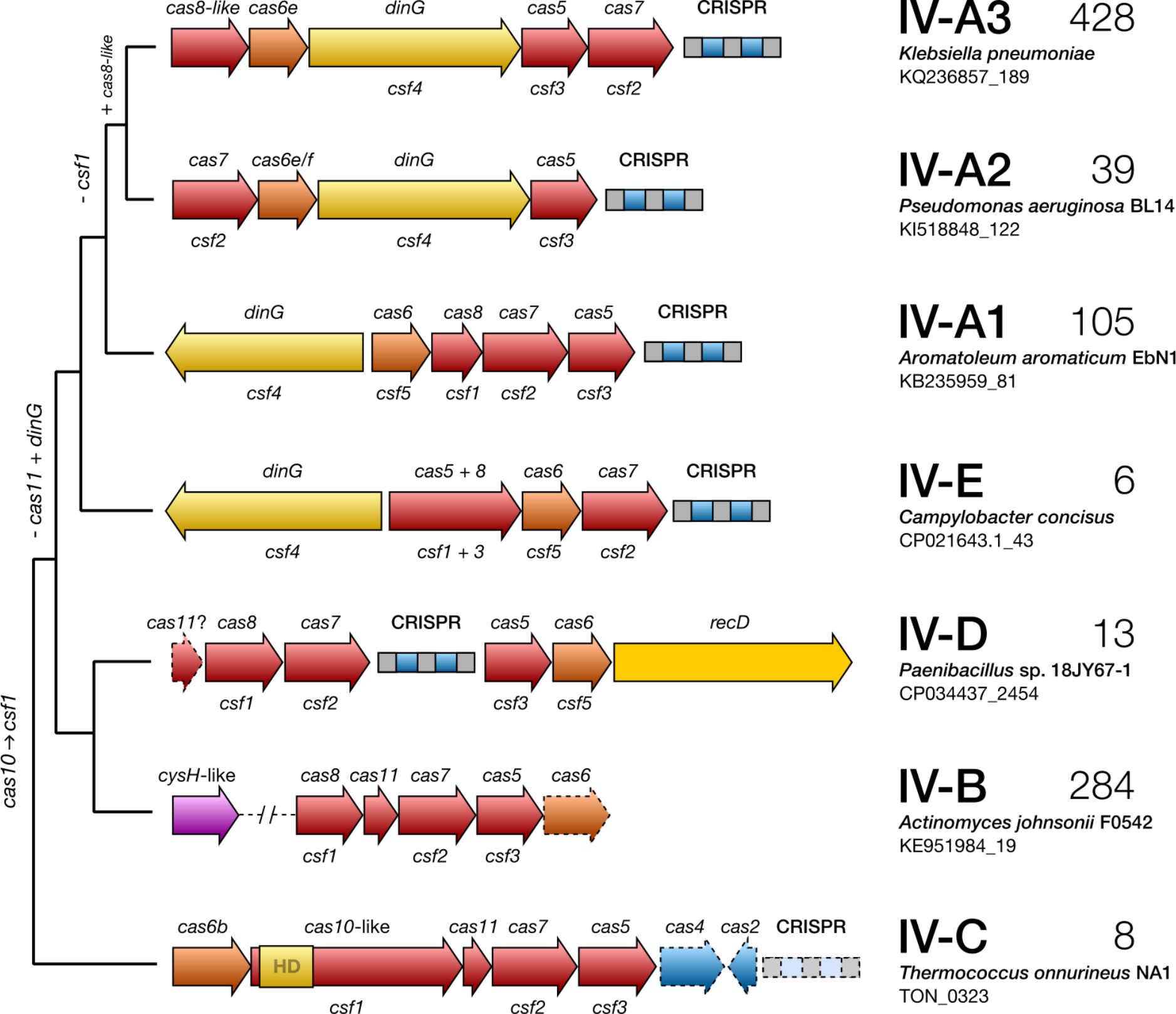
A proposed classification of type IV CRISPR-Cas systems based on their genome loci architectures and evolutionary relationships. Phylogenetic tree depicting the typical operon organization of the identified subtype IV loci. A selected representative locus is shown for each clade wherein genes are colour-coded and labelled according to the protein families they encode, using both the cas (upper) and csf (lower) nomenclatures. Genes or CRISPR arrays that are not invariably present are represented with dashed lines on the gene maps. The number of loci identified for each clade is given on the right. Hypothesised gene gain/loss events over the course of evolution are shown on the left.

Overall, subtypes IV-B and IV-A exhibit a high level of genetic diversity (Supplementary Fig. 3). Subtype IV-B is composed of several phylogenetically divergent clades, merged here because of their similar genetic architectures, and subtype IV-A spans three major groups, hitherto referred as IV-A variants 1, 2 and 3. Notably, although subtypes IV-A2 and IV-A3 are closely related, they primarily differ in the absence (IV-A2), or presence (IV-A3), of a gene in their cas operons. We infer this gene encodes a Cas8-like protein due to its shared features with other Cas8 components, such as similar size and a zinc finger domain (Supplementary Table 2). Since Cas8 proteins often show little or no significant sequence similarity, even within subtypes^1,16^ (e.g. subtype I-B), and because all three IV-A variants cluster as a monophyletic group showing comparable modular architectures (Fig. 1, Supplementary Fig. 3), we maintain them within the same subtype. Notably, the common exchange of functional modules^17^ between different CRISPR-Cas systems is particularly evident for IV-A2 and IV-A3, where Cas6 apparently has been recruited from subtype I-F and I-E systems, respectively (Fig. 1, Fig. 4b), highlighting a possible functional link between these subtypes.

Additionally, we identified a distinctive group of loci (named here subtype IV-D) which is unique in carrying a helicase of the RecD family in place of the archetypal DinG. This latter observation highlights the putatively central functional role of a dsDNA unwinding component in these systems. Moreover, while IV-B and IV-D appear to have diverged relatively recently, their classification into separate subtypes seems justified. Unlike subtype IV-D, IV-B loci are typically associated with a *cysH*-like gene (a member of the adenosine 5’-phosphosulfate reductase family^8,18^) and they do not encode a helicase, a Cas6 (with rare exceptions; Supplementary Data 1) or a CRISPR array. Finally, a few examples were found of an outgroup clade related to IV-A, labelled here as the putative subtype IV-E. Despite sharing similar modular architectures, their DinG components have diverged significantly (Supplementary Fig. 5) and the Csf1 of subtype IV-E is fused to Csf3, as revealed by HHpred searches (Supplementary Table 3).

### Type IV systems are widely distributed across taxa and diverse MGEs

Our taxonomic analysis reveals a widespread, yet heterogeneous, distribution of type IV loci across a variety of prokaryotic genome backgrounds and they were primarily predicted to be encoded by MGEs (Fig. 2, Supplementary Data 2). Subtypes IV-A and IV-B appear to be the most prevalent, contrasting with the sparse and relatively narrow taxonomic distribution of the other subtypes. While IV-A variants are mainly spread across proteobacterial plasmid-like conjugative elements, subtype IV-B is largely confined to predicted plasmids (and sometimes prophages) of Actinobacteria, and to a lesser extent Archaea, Firmicutes and Proteobacteria. The reduced group of subtype IV-C loci were found in Archaea, and no evidence for a preferential association with MGEs was found. Moreover, subtype IV-D occurs in some plasmids of Firmicutes and IV-E modules are present in Campylobacter and Bacteroides; some of the latter also residing in plasmid-like elements. Notably, in sharp contrast to the near-exclusive association of type IV systems with MGEs, we rarely found other CRISPR-Cas types to be encoded by plasmids or prophages, consistent with earlier reports and highlighting the uniqueness of type IV systems in this regard^1^.

**Fig. 2.**
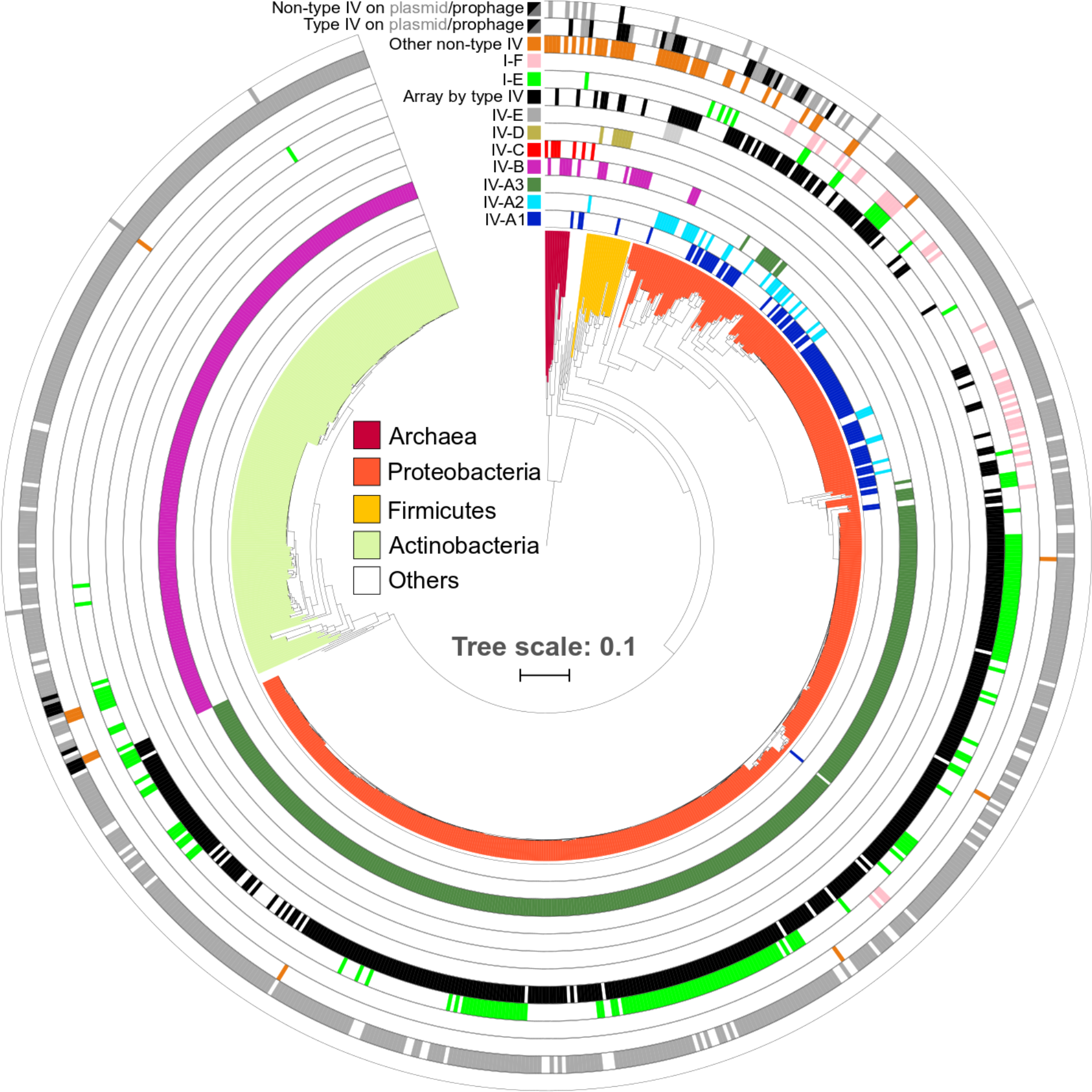
Distribution of type IV loci across prokaryotic taxa and MGE types. Phylogenetic tree based on 16S rRNA gene sequences of all bacteria and archaea that carry type IV CRISPR-Cas systems. Concentric rings denote the presence or absence of type IV and other co-occurring non-type IV CRISPR-Cas loci in the same genomes, colour-coded according to the subtype/variant to which they belong. All non-type IV systems, except I-E (light green) and I-F (pink), were merged into one lane (orange) for visualization purposes. Type IV effector cas operons for which an associated CRISPR array was detected are shown (black). Based on genomic context analyses (Methods), CRISPR-Cas systems predicted to be encoded by plasmid-like elements (grey) or (pro)phages/viruses (black) are shown (Supplementary Data 2), for both type IV and non-type IV loci (two outermost ring lanes). 758 (of 883) identified type IV loci are displayed on the tree; for the remainder no 16S rRNA gene sequence was found in the genome.

### Type IV spacer contents exhibit a strong bias towards plasmid protospacers

Statistical analyses of the distribution of spacer matches has proved a powerful tool for predicting functional properties of CRISPR-Cas systems and for understanding the ecology of the genomes carrying them^19,20^. Given that type IV loci are primarily harboured by plasmids, semi-independent entities with selective pressures differing from those of their hosts^21^, we sought to investigate whether type IV systems exhibit different targeting preferences from non-type IV systems. Therefore, we performed a comprehensive analysis of the spacer-protospacer matches for all the type IV-associated CRISPR arrays, and for the CRISPR arrays of all other identified non-type IV systems present in the host genomes.

Consistent with earlier results^1,8^, only a small fraction of spacers yielded significant matches: ~12% and ~7%, for type IV and non-type IV, respectively (Fig. 3a, Supplementary Table 4 and Supplementary Data 3 and 4), which reflects the current undersampling of the microbiome^22^. However, we observed that type IV systems displayed an exceptionally strong targeting bias towards plasmids, in contrast to the other co-occurring CRISPR-Cas systems (80% vs. 26%, respectively). Importantly, this trend was valid for all DinG associated type IV subtypes and variants, whereas the remaining subtypes did not yield sufficient data. On the other hand, non-type IV subtypes overall exhibited the previously reported strong preference for viral targets (Fig. 3a., Supplementary Table 4)^1,22^. Given that the type IV and non-type IV spacer contents investigated here originate from the same cellular environments, the results strongly underline an anti-plasmid function for type IV systems.

**Fig. 3.**
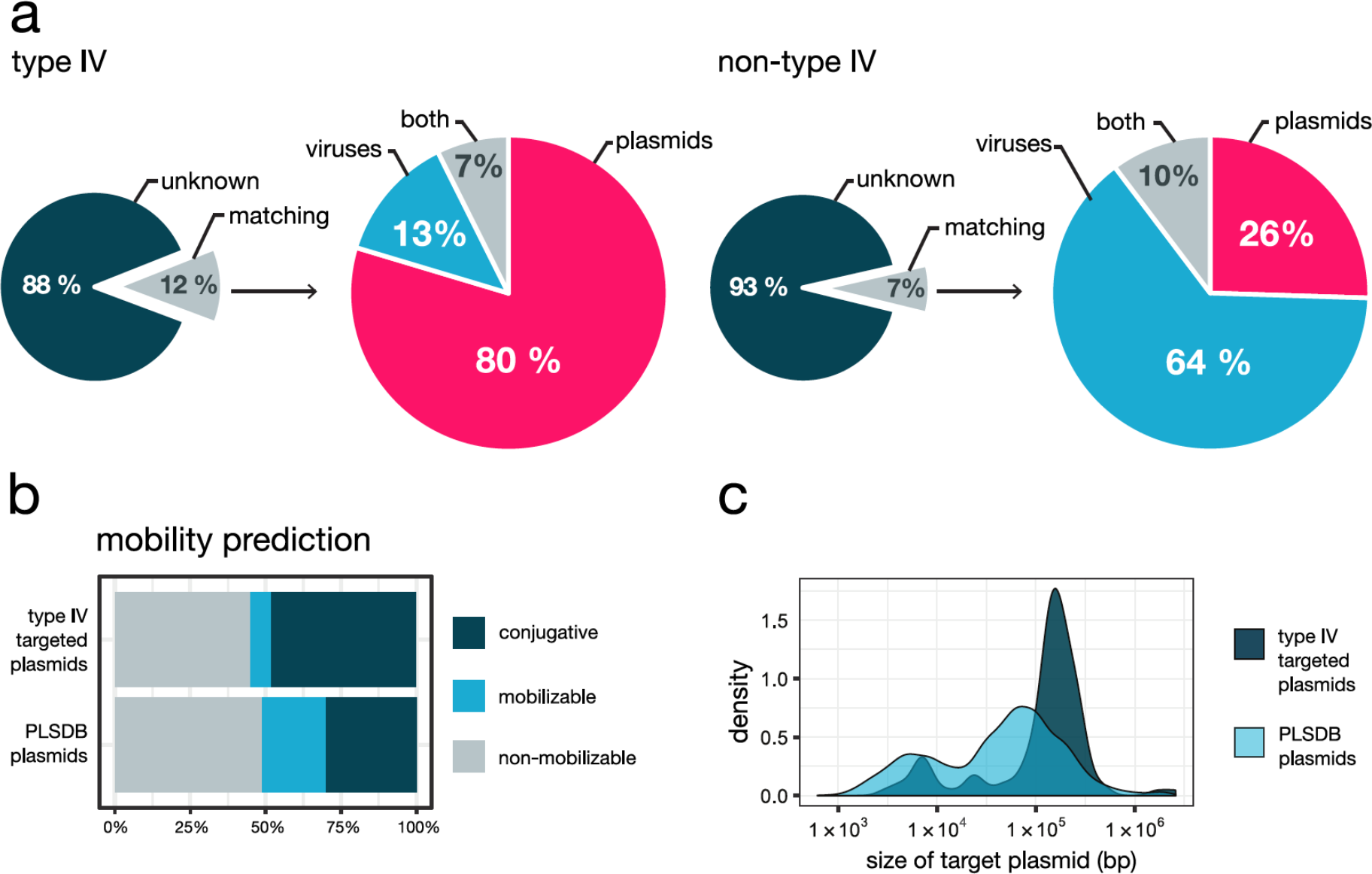
Spacers from type IV systems preferentially target plasmid-borne protospacers. **a.** Comparison of spacer-protospacer matches detected for type IV systems (left) and the co-occurring non-type IV systems (right). A more detailed breakdown, by CRISPR-Cas subtype/variant, is presented in Supplementary Table 4. **b.** Distribution of type IV spacer hits on plasmids as a function of predicted plasmid mobility. **c.** Size distribution of the targeted plasmids. The mobility prediction and size for the collection of PLSDB plasmids are displayed as a reference in both “b” and “c” plots.

**Fig. 4.**
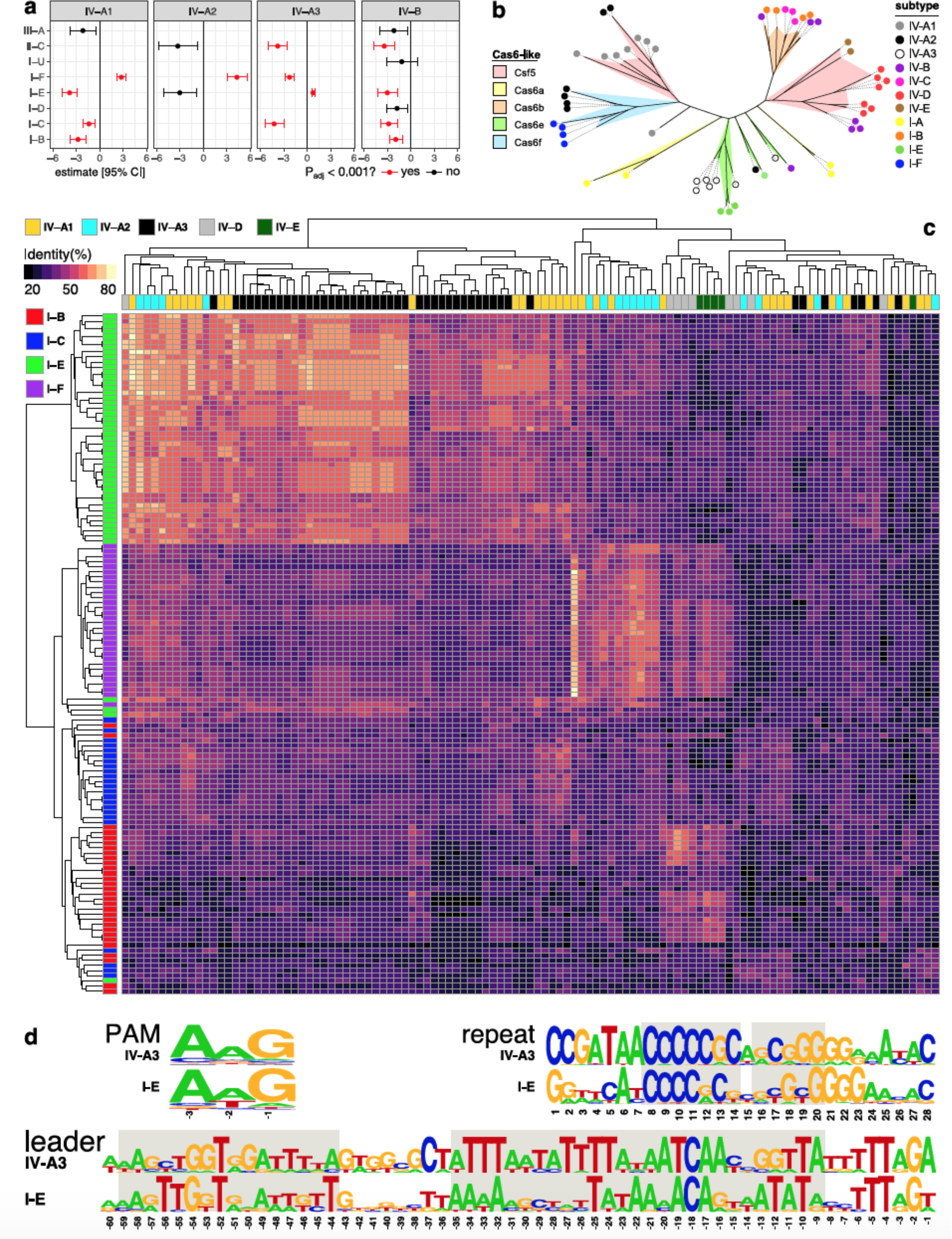
Interactions between type IV CRISPR-Cas systems and other co-encoded CRISPR-Cas systems in a host. **a.** Co-occurrence analysis between type IV and non-type IV systems. Estimates are from phylogenetic logistic regressions, with p-values fdr-adjusted. Only estimates with standard errors <10 are shown. **b.** Unrooted phylogenetic tree for Cas6/Csf3 built with representatives covering the diversity of type IV and type I subtypes/variants detected in this study. Each cluster is coloured according to the cas6-like family it corresponds to, and the coloured dot at the end of branches indicates the specific CRISPR-Cas subtype/variant encoding such a Cas6-like protein. **c.** Heat map depicting CRISPR repeat similarity of co-occurring CRISPR-Cas subtypes/variants clustered by average linkage hierarchical clustering. **d.** PAM, consensus CRISPR repeat and leader sequence logos for the positively correlated subtypes IV-A3 and I-E. The short semi-palindromic repeats at the centre of the consensus repeat that are used as anchor sequences by the Cas1-Cas2e complex are highlighted in grey, as well as the conserved leader sequences comprising the binding sites for the Cas1-Cas2e complex (left) and the IHF (right).

Next, in order to further explore the potential functional differences between type IV and non-type IV systems, we examined possible variations in their targeting preferences towards specific plasmid and viral gene families. Statistical analyses of the spacer match distributions revealed an enrichment of certain plasmid and virus-related genes, yet no consistent differences were observed between type IV and non-type IV targets (Supplementary Fig. 6 Supplementary Data 3 and 4). In agreement with previous reports^22^, all CRISPR-Cas types revealed a targeting preference for conserved, and frequently plasmid-borne, genes; e.g. conjugative transfer machinery genes (Supplementary Fig. 6). Although a similar pattern was observed for non-type IV viral gene matches, the corresponding analysis for type IV was inconclusive due to the low number of identified viral protospacers. Next, we investigated whether the plasmids targeted by type IV-derived spacers displayed any unifying biological features that could provide insights into the function of type IV systems. In general, we found that targeted plasmids tend to be relatively large (Fig. 3c., Targeted: 155 kbp, PLSDB: 53 kbp, median sizes, P<2.2*10^−16^, Kolmogorov–Smirnov test), irrespective of their predicted mobility (Supplementary Fig. 7), and there was a clear bias towards the targeting of conjugative plasmids (Type IV: 48%, PLSDB: 30%, Fig. 3b).

### Type IV associations with other CRISPR-Cas systems

The almost exclusive absence of adaptation module genes from type IV loci (Supplementary Data 5) raises the question as to the origin of CRISPR spacers. This aspect of type IV’s biology is especially puzzling given the observed variability in spacer content between related type IV CRISPR loci. Notably, spacer acquisition invariably requires Cas1 and Cas2, the most conserved components of all CRISPR-Cas systems^23^. This high conservation implies that there could be degrees of compatibility between adaptation modules of different CRISPR-Cas types. Therefore, we reasoned that type IV loci could exploit this functional redundancy by co-opting Cas1/Cas2 adaptation modules from other CRISPR-Cas systems that coexist intracellularly.

To explore this hypothesis, we first searched for evidence of positive correlations between different type IV subtypes/variants and other CRISPR-Cas systems present within the same hosts (Fig. 4a). Interestingly, significant positive correlations were found for subtypes IV-A1/2 and IV-A3, together with subtypes I-F and I-E, respectively (IV-A3 and I-E: P=3.7*10^−13^, IV-A2 and I-F: P=4.5*10^−10^, IV-A1 and I-F: P=3.7*10^−17^, fdr-adjusted P-values from phylogenetic logistic regression). We also found significant negative correlations between several subtypes, including IV-A1 with I-B, I-E and I-C, IV-A3 with I-C, I-F and II-C, and IV-B with I-B, I-C, I-E and II-C, which could be due, at least partly, to the targeting of type IV-carrying plasmids/MGEs by host-encoded CRISPR-Cas systems. Furthermore, co-clustering of CRISPR repeats demonstrated that type IV repeat sequences are similar to those from CRISPR loci with which they co-occur and/or correlate positively (IV-A1/2, IV-A3 and IV-D, with I-F, I-E and I-B, respectively) (Fig. 4c), strengthening the notion of a potential functional connection between type IV and other co-encoded CRISPR-Cas systems.

PAM (protospacer adjacent motif) recognition is essential for Cas1/Cas2-dependent spacer acquisition and self/non-self discrimination in most CRISPR-Cas systems^24–26^, yet such motifs have not yet been described for type IV systems. Therefore, we investigated whether PAMs could be identified and, if so, whether they are compatible with co-occurring non-type IV CRISPR-Cas systems. To test this, we predicted PAMs *in silico* by aligning protospacer flanking regions and putative PAM was identified for subtype IV-A3 (Fig. 4d). However, searches for other subtypes/variants were unsuccessful, likely due to the low number of spacer-protospacers matches (Supplementary Table 4). Importantly, the predicted subtype IV-A3 PAM (-AAG-) is identical to that of the positively correlating type I-E CRISPR-Cas system.

The higher numbers of detected subtype IV-A3 loci provided the basis for a case study involving more extensive comparative analyses. Alignments of consensus repeats of the positively correlating subtypes IV-A3 and I-E (Supplementary Fig. 8) revealed the previously described recognition sites for the Cas1-Cas2e adaptation machinery^27^ (Fig. 4d). In addition, multiple sequence alignments of the upstream regions from co-occurring IV-A3 and I-E CRISPR arrays (Supplementary Figs. 9a and 9b, respectively) showed similar conserved motifs in the leader region (Fig. 4d). Importantly, these conserved sequences comprise the binding sites for the Cas1-Cas2e complex and the integration host factor (IHF), both of which are essential for uptake of new spacers into leader-repeat junctions of type I-E arrays^28,27^. Next, we searched for evidence of preferential acquisition of spacers in the leader-end of IV-A3 CRISPR arrays, a phenomenon described for some CRISPR-Cas systems^29,30,31,32^. However, clustering of related IV-A3 CRISPR loci and a comparison of their spacer contents did not reveal any clear support for this (Supplementary Fig. 10). Finally, analysis of type IV-associated Cas6-like proteins yielded evidence for a polyphyletic origin, with independent acquisitions having occurred on multiple occasions (Fig. 4b). For example, IV-A3 and IV-A1/2 loci contain Cas6 variants that are more closely related to Cas6e and Cas6f, respectively, than to Csf5^33^, and co-occurring IV-C carries a Cas6b enzyme, further underlining the functional interrelations occurring between type IV and type I systems.

## Discussion

In addition to the recognised adaptive immune functions of CRISPR-Cas systems, there is increasing evidence that diverse MGEs, including phages, giant viruses and transposons, have co-opted these systems for alternative functions^34,7,35,36,37^. The discovery of type IV systems, and of their frequent encoding on plasmids, is also relatively recent^1^. To date, only two subtypes (IV-A and IV-B) are known and their biological functions and mechanisms of action remain obscure. In this work, we identify several novel type IV subtypes/variants and incorporate them into a revised type IV classification (Figure 1). In agreement with previous work, we found that the newly identified type IV loci are primarily encoded by prokaryotic MGEs, most of which are predicted to be plasmids (Figure 2). Notably, given the current limited sequence information covering the “dark matter” of the mobilome^22,35,36^, our findings likely underestimate the true diversity and distribution of these systems. Future comparative genomic characterizations will clearly benefit from including metagenomic sequence datasets and the continuing global effort to sample the meta-mobilome.

Our analyses suggest an origin of type IV systems from a type III-like ancestor in archaea (most similar to the subtype IV-C described in this work), comparable to the evolutionary pathway proposed for the emergence of type I CRISPR-Cas systems^15^. This is further supported by IV-C a) carrying a Csf2 with structural similarities to Cmr4/Csm3, the Cas7-like helical backbone subunit in subtypes III-A/B (Supplementary Fig. 11), b) being exceptional for type IV in carrying an HD domain on its Cas10-like protein instead of a helicase and Cas8/Csf1 (Fig. 1), c) being most commonly found in archaeal hyperthermophile genomes and d) being only intermittently associated with CRISPR arrays and type I systems (Supplementary Fig. 4), all of which are characteristic properties of type III systems. With regard to the evolution of the remaining type IV subtypes, a parsimonious scenario involves streamlining of the Cas10-like protein into Csf1 generating a IV-B-like ancestor that acquired a RecD helicase which led, in turn, to the evolution of subtype IV-D. In a separate branch such a IV-B-like ancestor is speculated to have lost Cas11 and acquired DinG leading to the evolution of subtypes IV-A and IV-E. The fusion between Csf3 and Csf1 in IV-E is consistent with the proximity of these proteins in class 1 effector complexes. As for IV-A, although the three variants are all closely related, IV-A2 appears to derive from IV-A1 after loss of Csf1, but retaining CRISPR-Cas functionality, while IV-A3 is a more recent variant of IV-A2 that seems to have gained a substitute for Csf1, the Cas8-like protein.

This evolutionary scenario is underpinned further by the overall taxonomic distribution of the identified type IV loci, ranging from the broad occurrence of IV-B, to derived variants such as IV-A3 being restricted to a few genera of Proteobacteria (Fig. 2). Interestingly, the emergence of IV-D from an IV-B-like ancestor pool may have occurred more than once, as evidenced by the paraphyly of IV-Ds (Supplementary Fig. 3). Subtypes IV-D, IV-A and IV-E are CRISPR array-associated, unlike IV-B which is more diverse and CysH associated. This likely reflects convergent evolution where type IV systems initiated as CRISPR-Cas immune system and then evolved an altered functionality before reverting back into CRISPR-Cas systems via lateral acquisition of a DNA helicase.

Contrary to the strong viral targeting preference of all other known CRISPR-Cas types, our work reveals that type IV systems exhibit an exceptional targeting bias towards plasmid-like elements (Fig. 3a, Supplementary Table 4). Intriguingly, this bias was not detected in earlier studies that employed lower numbers of non-redundant spacers and were primarily centred around the matching of spacers against (pro)virus databases^7,9,22,38^. Our additional matching of spacers against PLSDB, a comprehensive database of >16,000 curated plasmid genomes^39^, was key in determining the plasmid targeting bias. Importantly, since our analysis of the spacer contents from other non-type IV CRISPR-Cas systems coexisting intracellularly with type IV loci clearly yielded the established bias towards viral targets (Fig. 3a, Supplementary Table 4), and both analyses were done matching spacers against the same databases, we conclude that the reported type IV plasmid bias cannot be an artefact.

Interestingly, a significant enrichment of certain targeted gene families was observed (Fig. 3b), particularly those encoding components of complex molecular machineries including the conjugative transfer apparatus (Supplementary Fig. 6, Supplementary Data 3). An explanation for this phenomenon is that conserved genes are less prone to mutational escape from CRISPR-Cas targeting, and thus lead to a positive selection of their cognate spacers over time^40^. Moreover, spacer retention may also be further enhanced when a targeted gene is shared by distinct MGEs, as also occurs, for example, with conjugative transfer genes.

The finding that type IV systems are carried by plasmid-like elements that primarily target other plasmids leads to the basic question as to how and why an anti-plasmid bias emerged. Our results indicate that type IV systems may have evolved to target plasmid-like elements more effectively than, for example, phages/viruses, although the mechanistic basis of such a bias remains unclear. Moreover, certain plasmids may provide strong competition for type IV CRISPR-Cas-carrying plasmids, leading to the selection of spacers against the former plasmids over time. Whereas phages/viruses can interfere with plasmid survival by killing the host, cells already carry potent defence systems against these fatal intruders^41^. Therefore, plasmids may be more strongly challenged by other intracellular plasmid-like elements which, while not being especially detrimental to the host, may compete directly for common cellular resources^42,43^. The latter argument receives support from the accepted community ecology view that similar entities compete more strongly for overlapping niches and resources^44,45,46^. Notably, recent work has proposed that many (pro)phages readily engage in similar CRISPR-based inter-virus warfare dynamics, utilizing “mini-arrays” with spacers targeting viruses to prevent host superinfection^7^. In summary, our results imply that plasmid-like elements leverage type IV systems to eliminate other plasmids with similar properties and lifestyles, in order to monopolize the host environment.

In addition, our findings reveal an apparent functional cross-talk between type IV modules and other co-occurring CRISPR-Cas systems within a host, thereby providing a credible explanation for the minimal nature of type IV systems. Not only did some type IV subtypes correlate positively with specific type I subtypes (Fig 4a) but there were also additional parallels between some co-occurring pairs: PAM sequence sharing, high CRISPR repeat sequence similarity and a high similarity between the Cas6 processing enzymes (Fig 4d,c,b, respectively). Noteworthy, future experimental work is required to both validate the predicted IV-A3 PAM and establish whether the large subunit Csf1 facilitates PAM recognition on nucleic acid targets as occurs for its homolog Cas8 in type I systems^47,48^. Furthermore, we also found shared conserved CRISPR leader motifs for the binding of the Cas1/2e adaptation machinery and IHF between co-occurring IV-A3 and I-E subtypes (Fig. 4d). Although all these results are consistent with the inference that type IV systems can rely on the Cas1-Cas2 adaptation machinery from co-occurring “helper” type I systems, such co-functionality requires experimental validation. Nevertheless, this hypothesis receives support from the numerous accounts of type III systems lacking Cas1 and Cas2 which utilise CRISPR-arrays maintained by adaptation modules from neighbouring type I systems^49,50^, and is further reinforced by the evolutionary links demonstrated here between type IV and type III systems.

Nevertheless, alternative spacer acquisition strategies cannot be ruled out. These include, for example, the mechanism proposed for viral-derived orphan mini-arrays, where recombination with host CRISPRs seems most likely^7^. Consistent with the latter hypothesis, we observe examples of spacer rearrangements between related IV-A3 CRISPR loci (Supplementary Fig. 10). Interestingly, most type IV systems carry a Cas6-like component, suggesting that specific pre-crRNA processing may be necessary for exclusive crRNA coupling with type IV effector complexes. This extra level of specificity and the stark contrast in spacer targets between positively correlating type I and type IV systems indicates that, although there are functional ties at the adaptation stage, the crRNA utilisation stage operates independently. Moreover, type IV systems may benefit from carrying their own Cas6 component by ensuring control over crRNA processing, especially in cells where no host-derived Cas6 is available.

Although elucidation of the specific targeting mechanism of type IV systems requires an experimental approach, it is likely that the associated helicase (DinG/RecD) is involved. Although our exhaustive analyses did not locate a nucleolytic active site in these enzymes, the presence of a cryptic nuclease domain is possible. In such a case, RecD/DinG could function mechanistically similarly to Cas3, the helicase-nuclease effector component of type I CRISPR-Cas systems^51,52^. Interestingly, similarly to Cas3, some non-type IV-associated DinG helicases have evolved 3′→5′ exonuclease activity^7,53^. However, even in the absence of a nuclease component, type IV systems could still co-opt host-encoded restriction enzymes to cleave their targets, possibly by rendering them susceptible to degradation upon dsDNA unwinding. In support of this, type III systems have recently been shown to utilise host degradosome nucleases to ensure successful interference of diverse MGEs^54^. Intriguingly, chromosomally derived RecD homologs are known to take part in the RecBCD complex (exonuclease V) which, in addition to playing a role in DNA repair, carries out defence functions through the degradation of invading genetic elements^55^.

Binding of type IV effector complexes to DNA could also destabilise the target, especially if it constitutes a rapidly replicating element. The consequences of replication fork collisions with protein-nucleic acid complexes (e.g. the transcription machinery) on genome integrity are well documented and can include replication fork arrest, premature transcription termination, and double-strand DNA breaks^56^. Notably, these physical conflicts are also known to destabilise plasmids, eventually leading to their extinction from within cell lineages^56,57^. The latter explanation is compatible with the hypothesis that type IV systems function similarly to the artificially developed catalytically dead CRISPR-Cas systems, which bind DNA targets but lack cleavage activity (eg. dCas9)^58^. These so-called CRISPR interference (CRISPRi) systems, silence the expression of targeted genes by blocking transcription factor binding or RNA polymerase elongation^58^. Moreover, type IV-mediated gene silencing could serve purposes beyond plasmid-plasmid warfare, such as altering host expression profiles to enhance plasmid propagation and/or stabilise maintenance, all piracy practices which plasmids are known to invoke via diverse mechanisms^59,60,61^. In the context of CRISPRi functionality, for which R-loop formation between the crRNA and the DNA target is key, the common association of DinG with type IV loci appears paradoxical, as it is well documented that the substrate for this helicase are R-loops that block replication fork advancement ^62,63,64^. Thus, it is tentative to speculate about the potential regulatory or antagonistic role of the helicase component in the removal of type IV crRNA-DNA hybrids, although the purpose of such a function remains unclear. Interestingly, *dinG* sometimes appears in the opposite orientation to the other genes in type IV loci (Fig. 1), consistent with the notion that its expression might be controlled independently.

Subtype IV-B systems constitute the most reduced and enigmatic version of type IV systems, lacking identifiable CRISPR arrays, Cas6, and a helicase component. This exceptional combination of features led to the proposition that it performs a different function from the other type IV systems e.g. similar to transposon-encoded CRISPR-Cas systems^7,61,35^. Because type IV-B systems encode all the necessary components to generate a Cascade-like surveillance complex (Csf1,Csf2,Csf3), we hypothesized that it could accommodate pre-processed crRNAs originating from other co-occurring CRISPR-Cas systems. However, we found no evidence of neighbouring CRISPR arrays, mini-arrays, SRUs^7^ or of palindromic sequences that could yield the characteristic stem-loop secondary structures of crRNAs. Interestingly, our data revealed significant negative correlations of IV-B with the presence of all other CRISPR-Cas systems in the hosts (Figure 4a). Taken together, it seems plausible that these systems could have been repurposed by plasmids/phages to bind and neutralise crRNAs that become available, thereby antagonising other CRISPR-Cas functions in the intracellular milieu. Nonetheless, the complexity of such an anti-CRISPR (Acr) mechanism would greatly contrast that of all other Acrs described to date ^65^, thus rendering this explanation unlikely. The key to deciphering the function of subtype IV-B possibly resides in its obscure, nearly invariant, genomic association with *cysH,* a protein which seems to have co-diversified with this subtype (Supplementary Fig. 12). Since *cysH* belongs to the phosphoadenosine phosphosulfate reductase family, to which DNA phosphorothioate modification enzymes also belong, these systems could be involved in epigenetic silencing, or either linked to or antagonising, related RM functions ^7,65,66^.

Collectively, our results provide further evidence of the strong dynamic pairing between CRISPR-Cas systems and MGEs. This complex co-evolutionary interrelation fits the described “guns for hire” paradigm, where CRISPR-Cas components are recurrently co-opted by different genetic entities for myriad defence and offence functions^6^. Noteworthy, repurposing the power and programmability of type IV systems for controlling plasmid propagation presents promising biotechnological applications, particularly in the face of the current growing concerns regarding the spread of virulence and antibiotic resistance determinants within and between microbiomes^67,68^. Indeed, as the mysteries surrounding the biology of type IV systems continue to be unveiled further opportunities will arise for expanding the CRISPR-Cas molecular toolbox.

## Methods

### Detection, clustering and classification of type IV modules

Bacterial and archaeal complete and draft genomes were obtained from genbank and scanned with the TIGR03115 Csf2 model^69^ using HMMER3^70^. Protein sequences from two genes upstream and downstream of the detected *csf2* gene along with the Csf2 sequence itself were pooled and subjected to an all-against-all sequence comparison using FASTA^71^. A neighbour-joining tree was constructed using distances derived from the aggregate similarities between each module pair using a previously described method^16^. The tree was used to pick diverse representative type IV systems, which were then annotated manually using PSI-BLAST^11^ searches. Hits on type III CRISPR-Cas systems were purged from the Csf2 tree. Following manual annotation, the protein sequences from the refined representative modules were pooled for another all-against-all sequence comparison. Protein sequences were clustered using the method previously described^72^ and another aggregate module similarity tree was built for the refined representative type IV module set. The tree was overlaid with gene maps of the type IV modules marked with the obtained protein clustering information (Supplementary Fig. 2). This was used for devising the subtypes (Figure 1 and Supplementary Fig. 3).

### Spacer-protospacer match analysis

CRISPR arrays were detected with CRISPRCasFinder (4.2.17,^73^) and matched to a Type IV module if any predicted operon was within a 10kbp radius (distance to first gene in the operon, this cut-off was based on Supplementary Fig. 13). Non-type IV CRISPR-Cas systems were also detected with CRISPRCasFinder in the same genome assemblies where type IV systems were found, and typing was manually corrected when necessary. Arrays were matched to an operon if it was within 10kbp (distance to first gene in the operon, see Supplementary Fig. 13). Phage genomes were obtained from the April 2019 version of the millardlab.org phage database (http://millardlab.org/bioinformatics/bacteriophage-genomes/), and plasmid sequences were obtained from the PLSDB database (2019_03_05,^39^). In order to rule out false positive matches to conserved spacers within undetected arrays on plasmids, the putative arrays in the plasmid database were masked when detected via CRISPRCasFinder^73,74^, CRISPRdetect (2.2)^73,74^ and CRT (1.2)^75^. Furthermore, all unique repeats pertaining to arrays from the initial CRISPRCasFinder search were aligned (blastn -task blastn-short,^76^) against the masked PLSDB database, and putative arrays were defined if two or more matches (E-value < 0.1) were found within 100bp, and these regions were masked as well. Spacers from the initial CRISPRCasFinder search were collapsed into a unique spacer set using cd-hit-est^77^ in order to avoid overrepresented spacers from sequencing bias. The unique spacer set was aligned against the masked plasmid and phage databases using FASTA^71^ with an e-value cut-off of 0.05.

### Targeted gene enrichment analysis

Enrichment in spacer targeting of certain functions was done by first predicting ORFs in all plasmid and phage genomes using Prodigal^78^, and then clustering genes using the protein clustering algorithm previously described^72^. The observed number of matches to each gene cluster was compared to 10^5^ simulations of random draws from a binomial distribution with size *n* equal to the number of genes in the gene cluster and the probability 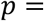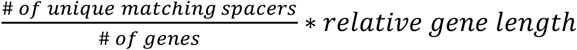, where the relative gene length was defined by dividing the length of each gene by the median gene length, and then finding the average length for each gene cluster. The above simulation only counted each spacer once, however, spacers usually match multiple genes in the same gene cluster. Therefore, each simulated match was multiplied by a random draw of the observed number of genes matched by a spacer matching that gene cluster.

### PAM identification

From the 1016 CRISPR arrays detected in Type IV containing complete and draft genomes, the consensus repeat for each array was aligned against corresponding consensus repeats for all other arrays using needleall^79^. Consensus repeats that differed from each other by more than two mismatches were assigned to separate repeat clusters, resulting in 171 repeat clusters in total. The previous unique spacer matching output from FASTA was surveyed for protospacers pertaining to each of the 171 repeat clusters separately. Spacers matching several phages and plasmids in the database were only counted once to circumvent sequencing bias in the database. Logo plots were drawn from the ten nucleotides immediately flanking each side of each unique protospacer. Protospacers with alignment lengths smaller than the total spacer length had their coordinates adjusted so all flanks within a repeat cluster were properly aligned.

### CRISPR-Cas subtype co-occurrence analysis

Co-occurrence between type IV and non-type IV subtypes was analysed with phylogenetic logistic regression (phyloglm, maximum penalized likelihood estimation^71,80^, with the non-type IV occurrence as the response and the type IV occurrence as the predictor. Besides the genomes with type IV systems, we supplemented the analysis with all complete genomes with at least one non-type IV operon (as defined by CRISPRCasFinder). The phylogenetic tree was based on 16S rRNA gene sequences detected by Barrnap^81,82^, aligned with mafft 7.307^83^, and tree made with FastTree2^81^, and was rooted by the archaeal clade. Edges of length zero were rescaled to the shortest non-zero branch length. Furthermore, outlier branches were pruned by removing tips for which the maximum phylogenetic variance-covariance was above 2. Only non-type IV subtypes found in at least 100 genomes were included, and only the four most prevalent type IV subtypes were included. P-values were fdr-adjusted with the Benjamini-Hochberg method ^84^.

### CRISPR repeat heatmap

All unique CRISPR consensus repeats were aligned with the pairwise2 module from Biopython 1.73^85^. Repeats were globally aligned with globalxs with both open gap and extend gap penalties of 3, and no end gap penalties. Alignments were done on both strands, and the highest identity was used.

### Leader sequence analysis

Multiple sequence alignments were performed with the upstream regions of a series of representatives of co-occurring IV-A3 and I-E CRISPR arrays using MUSCLE^86^. Alignments were analysed and visually displayed using Jalview ^85,87^. The corresponding leader sequence conservation profiles were generated using WebLogo 3^88^.

### Plasmid mobility prediction

The mobility of all plasmids (conjugative, mobilizable or non-mobilizable) in PLSDB was predicted with mobtyper^89^ with an E-value cut-off of 1e-10. For calculating whether certain mobility types were enriched in targeted plasmids, the number of matches were scaled such that the sum for each spacer was 1, which ensured that each spacer only counted once, no matter how many matches it had.

### Plasmid/prophage prediction

To predict whether the CRISPR-Cas operons were located on chromosomes or MGEs, we used an iterative heuristic; first, contigs from complete genomes were annotated as plasmids or chromosomes as described in the NCBI name. Second, for draft genomes PlasFlow^90^ was used to detect contigs that were putatively part of plasmids. Third, VirSorter^90,91^ was used to predict the presence of prophages, and all operons within a category 1, 2, 4, or 5 region were classified as putative prophages.

### Protein structure prediction

Protein homology models were generated with the Phyre2 protein structure prediction server (intensive mode)^92^. Superimposition of protein structures were generated by the PyMol molecular visualization software^93^.

## Acknowledgements

The authors would like to thank Jonas Stenløkke Madsen and Joseph Nesme for helpful discussions and useful comments. R.P.-R is supported by DFF Independent Research Fund Denmark, J.R. is supported by the Novo Nordisk Foundation, S.J.S. is supported by Lundbeckfonden, L.R is supported by the DFG SPP2141 and Heisenberg Programme and S.A.S is supported by grant A6291 from the Capital Region of Denmark.

## Author contributions

Conceptualization, R.P.-R., S.A.S., D.M.-M., and J.R.; Direction and planning, R.P.-R., and S.A.S. with support from S.J.S., Investigation and data analysis, R.P.-R., J.R., S.A.S., and D.M.-M., Writing, R.P.-R. with support from S.A.S., J.R., D.M.-M. and R.A.G., and in consultation with S.J.S. and L.R.

