## Supplementary Figures and Tables for "Type IV CRISPR-Cas systems are highly diverse and involved in competition between plasmids"

**Supplementary Fig. 1.** Maximum likelihood phylogeny of Csf2 (Cas7-like) from the collection of type IV-systems found in the study. (attached separately due to size constraints)

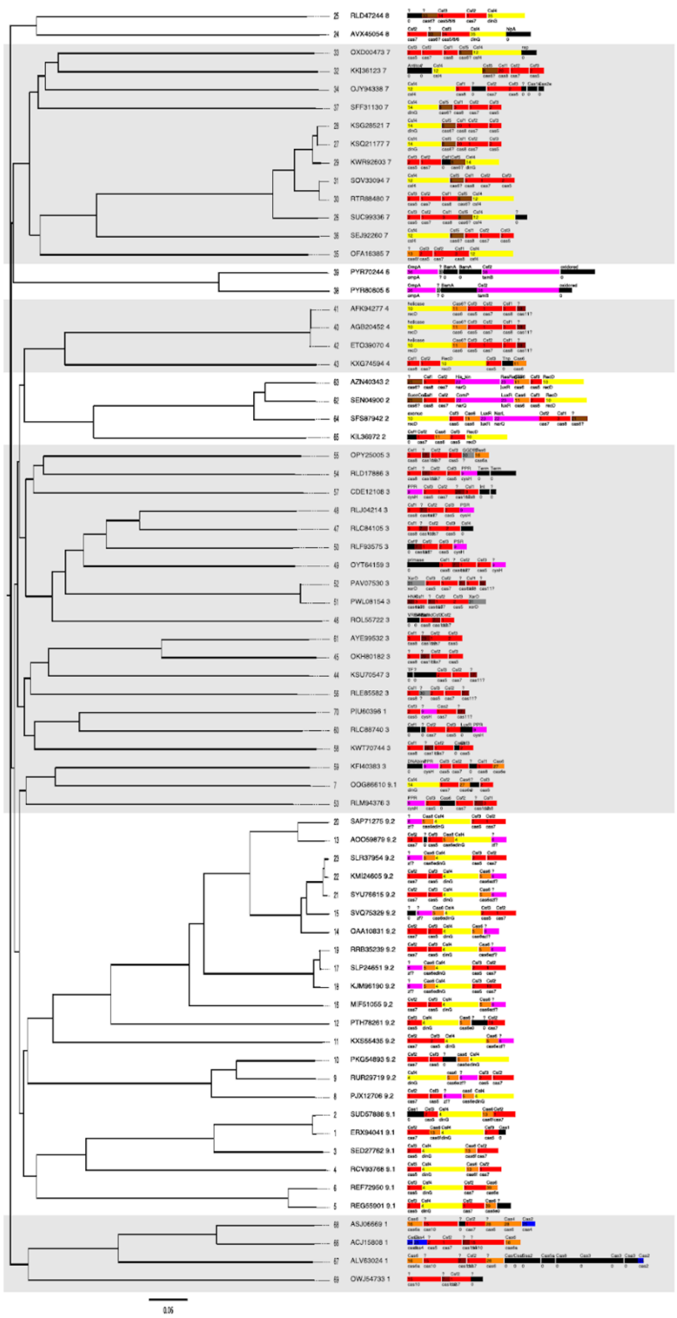

**Supplementary Fig. 2.** Full tree generated from 69 selected representatives of type IV loci that is based on aggregate protein similarity between all gene products. Gene products were assigned to orthologous gene clusters (numbered 1 through 34) using protein clustering and annotated based on profile-profile alignments against protein family databases (Methods) and PSI-BLAST. Core CASCADE subunits are colour-coded red, optional subunits dark red, Cas6 orange, helicases yellow, adaptation genes blue, accessory genes purple, putative Cas6s brown, and unknown conserved genes gray.

**Supplementary Fig. 3.** Phylogenetic tree of type IV and other co-occurring non-type IV CRISPR-Cas loci. (attached separately due to size constraints)

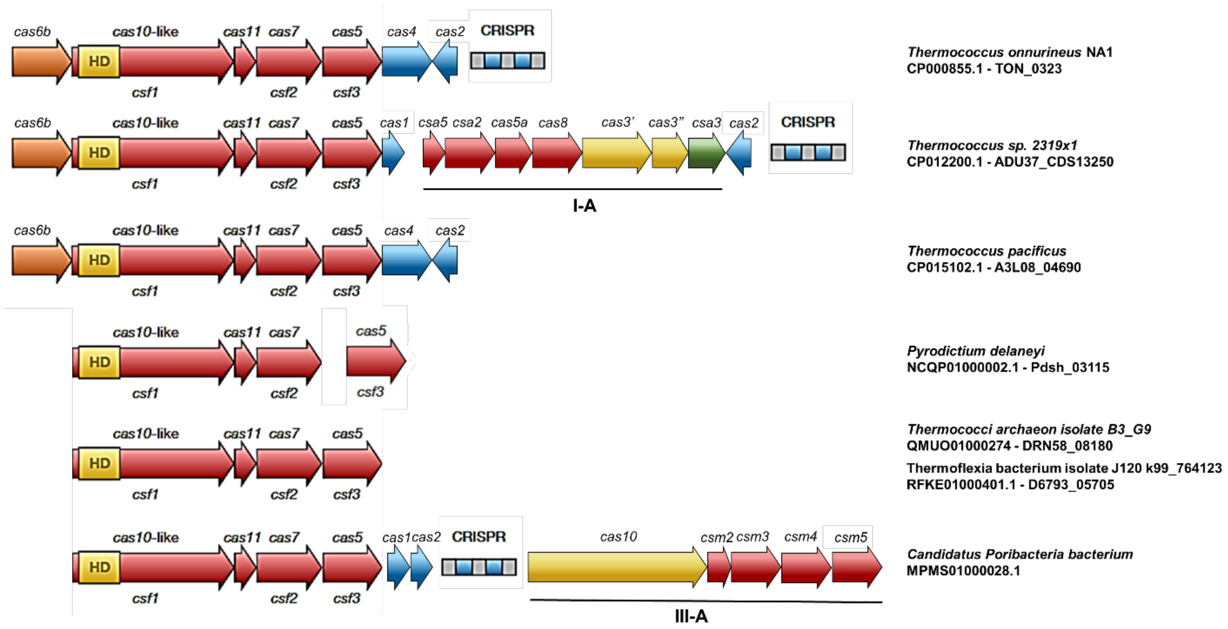


**Supplementary Fig. 4.** Subtype IV-C modules and associated CRISPR-Cas systems. Genes are colour-coded and labeled according to the protein families they encode, using both the cas (above) and csf (below) nomenclatures. Each loci is labelled in the left with the organism where is found as well as its accession number and the corresponding *csf2* gene ID.

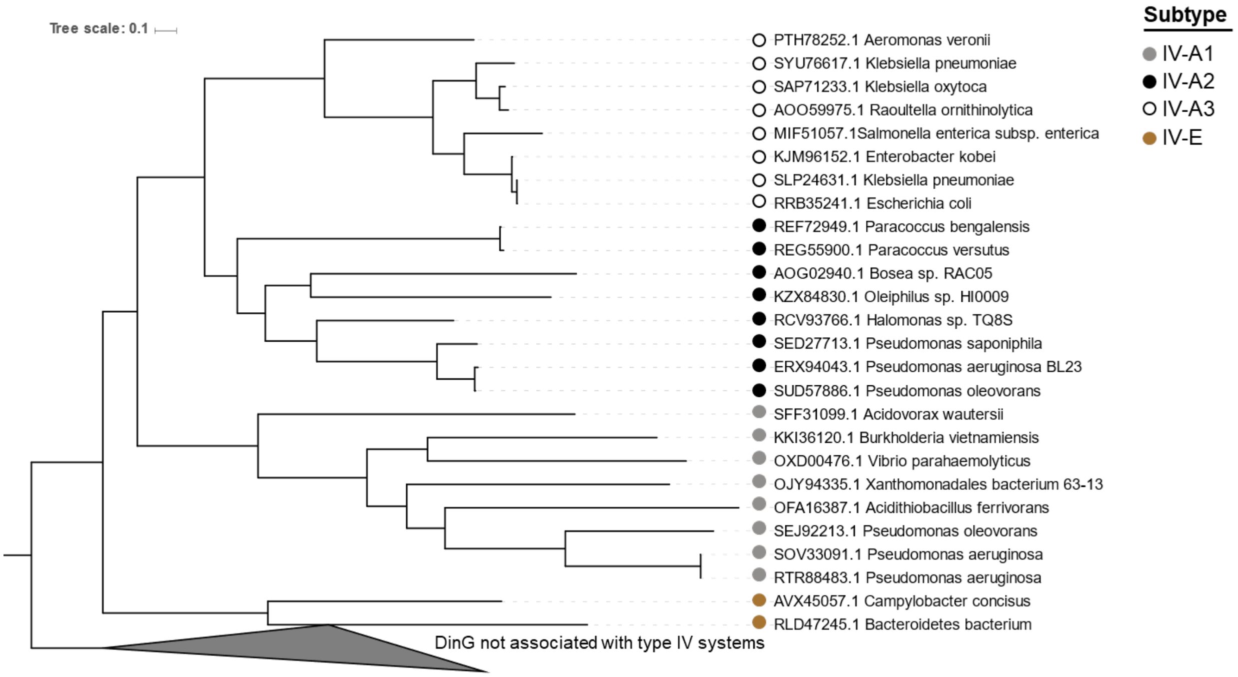


**Supplementary Fig. 5.** Phylogeny of helicases DinG associated with type IV systems, constructed with a subset of representatives from all type IV subtypes where one of the proteins was present. Non-CRISPR-Cas-associated DinGs were used to root the maximum likelihood tree. Branches are colour-coded according to the identity of the type IV subtype/variant, as illustrated in the figure.

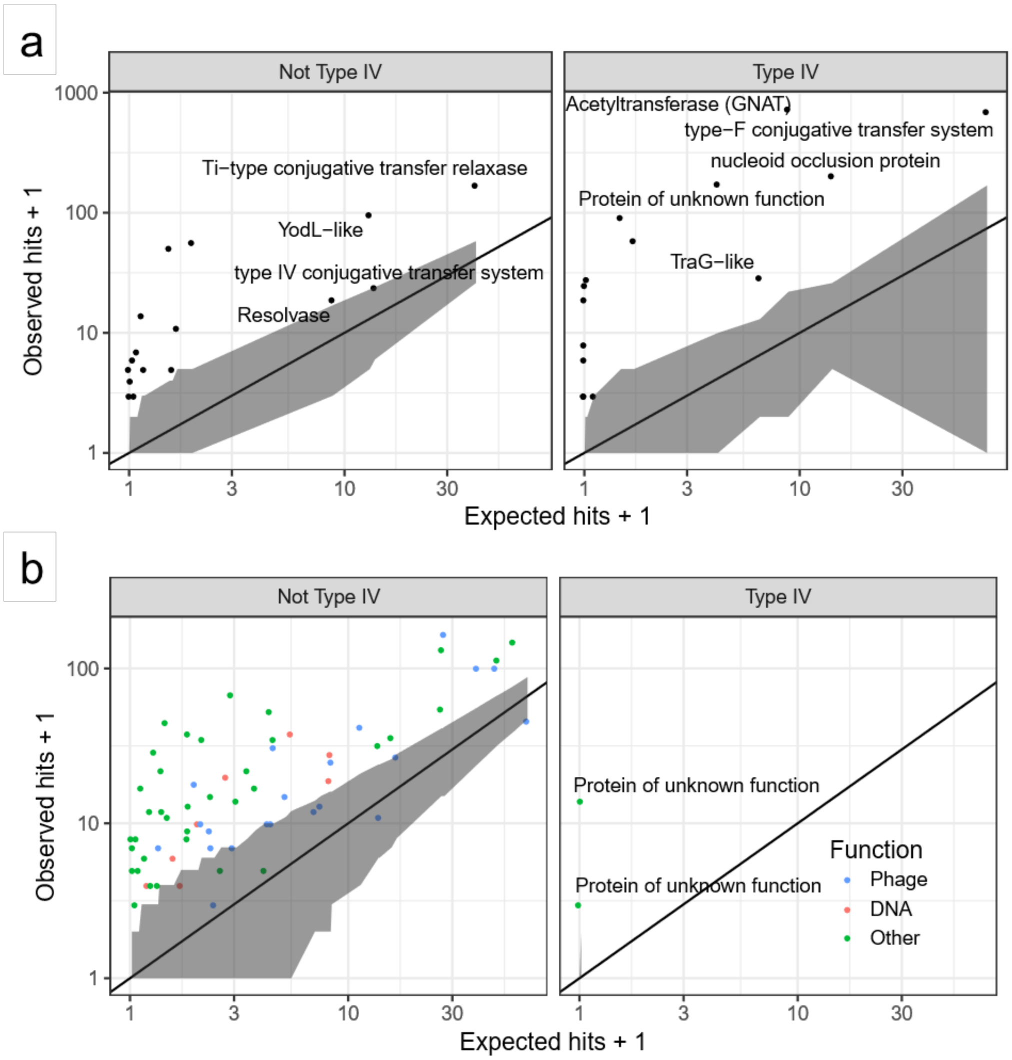


**Supplementary Figure 6.** Observed vs expected plasmid (a) and phage (b) spacer hits for all gene families targeted by type IV spacers (right) and non-type IV spacers (left). The expected number of hits were calculated by simulating random spacer-protospacer matches taking into account the size of gene clusters, length of genes, and the conservation of genes within a gene cluster (See Methods).

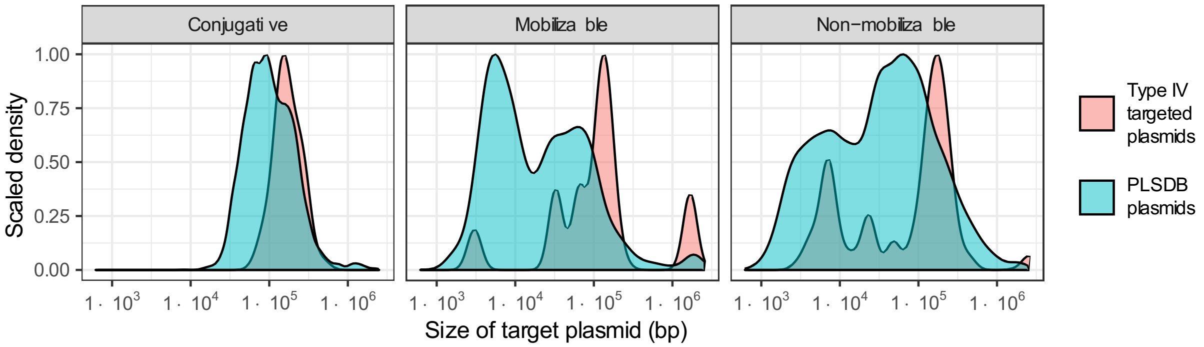


**Supplementary Fig. 7.** Size distribution of the plasmids targeted by spacers associated with type IV (red) CRISPR-Cas systems identified in this work, separated according to the predicted plasmid mobility type: conjugative, mobilizable or non-mobilizable. Size distributions for all plasmids deposited in PLSDB[40] are plotted as a reference (blue).

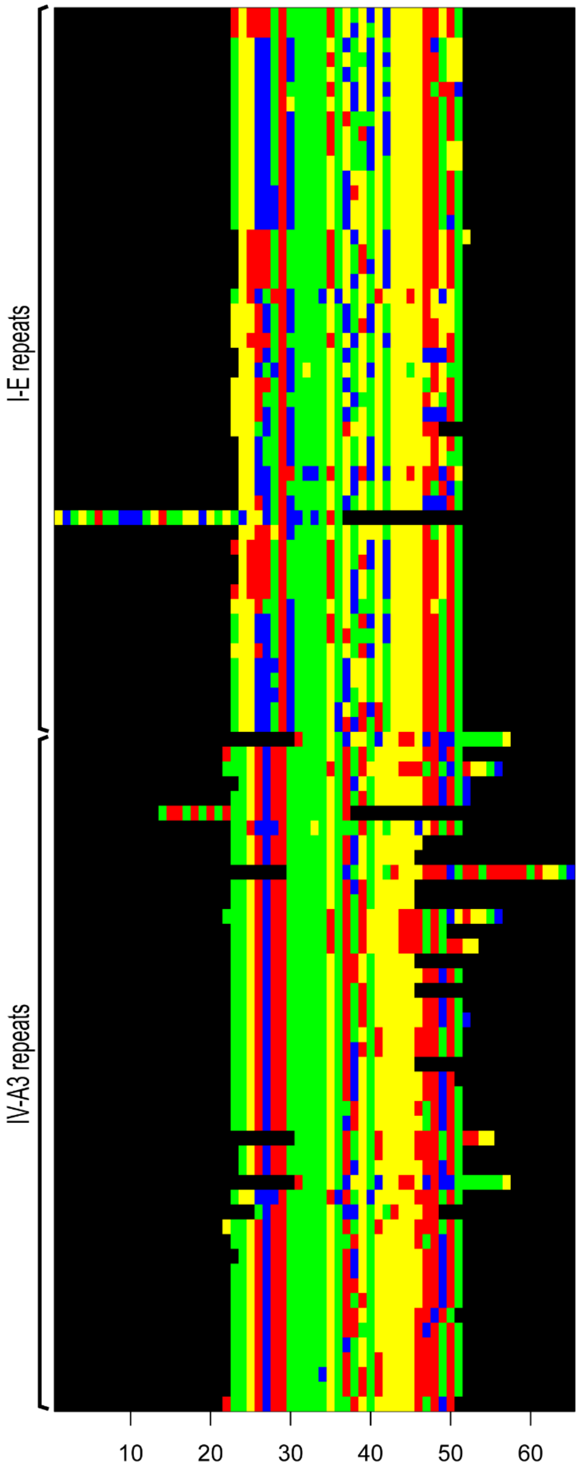


**Supplementary Fig. 8.** Alignment of CRISPR repeats of subtypes IV-A3 (bottom) and I-E (top) (mafft 7.307, with --op 3 --adjustdirectionaccurately). Each nucleotide is colour-coded: C - green, G - yellow, A - red and T - blue.

**a**

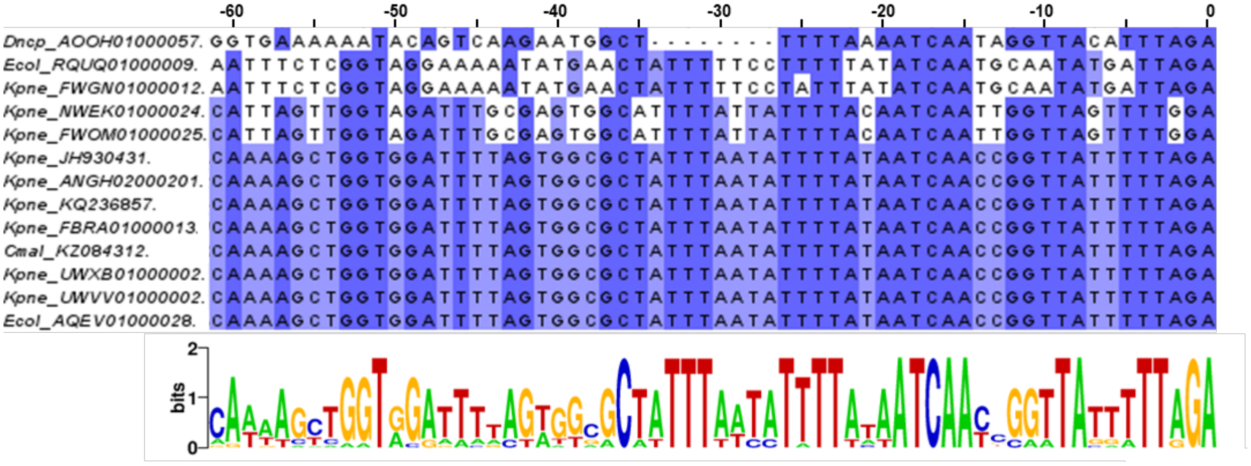


**b**

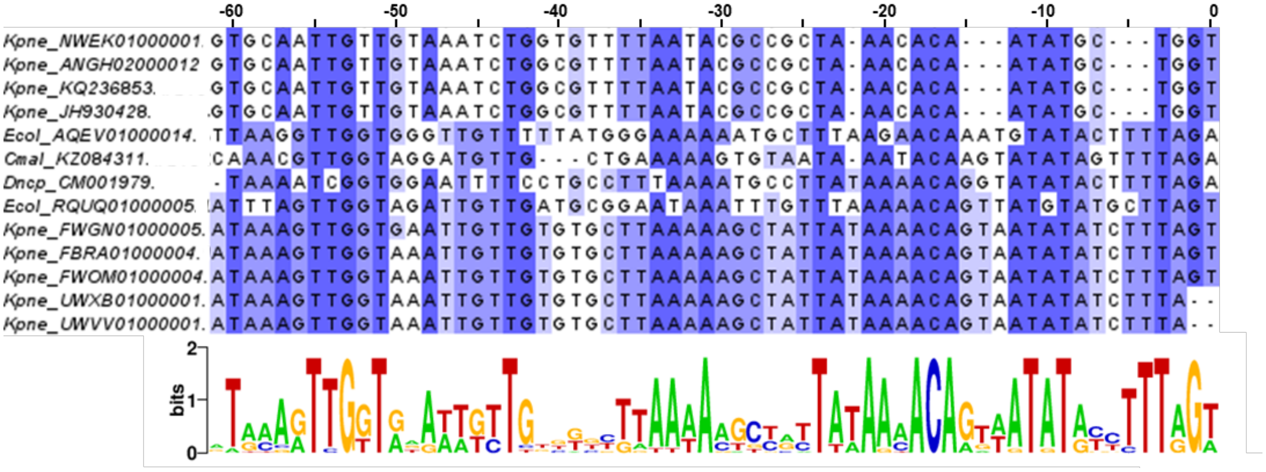


**Supplementary Fig. 9.** Multiple sequence alignments of the leader regions from co-occurring (**a**) IV-A3 and (**b**) I-E CRISPR arrays that originate from 13 arrays deriving from the IV-A3 variant cluster (Supplementary Fig. 4). Labels above the sequences denote positions relative to the leader-repeat junction. Residues in the alignment are highlighted in blue to reflect the extent of conservation using Jalview, where darker blue indicates a higher level of conservation. The leader conservation profile depicted in the lower section was generated using WebLogo3.

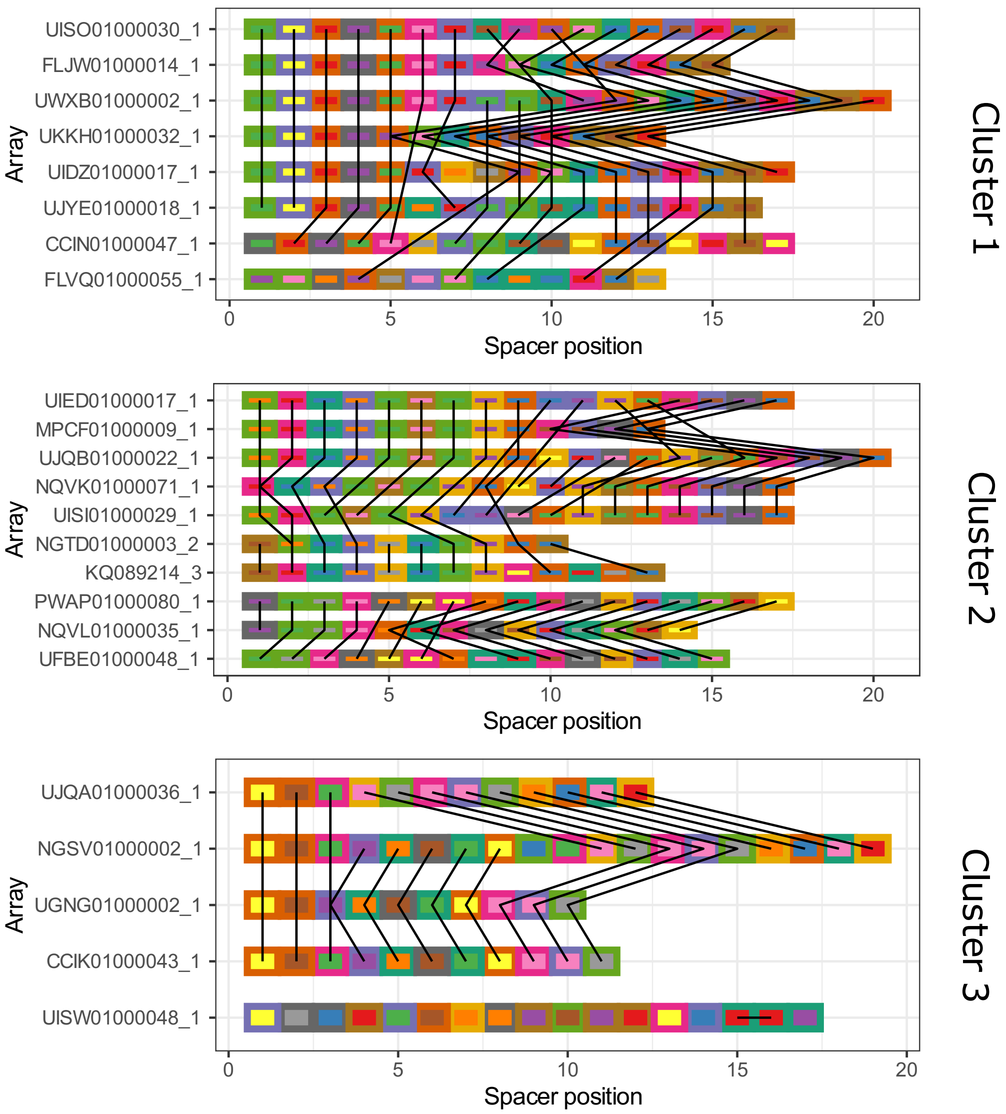


**Supplementary Fig. 10.** Spacer comparisons of similar arrays. IV-A3 CRISPR arrays were first reduced to a non-redundant set (95% similarity) and then clustered if they showed >80% identity across the array. The plot shows the three largest IV-A3 array clusters found. For each array cluster, similar spacers (>95% identical across >95% of length) are connected with lines and colour-coded. Clustering was performed with cd-hit.

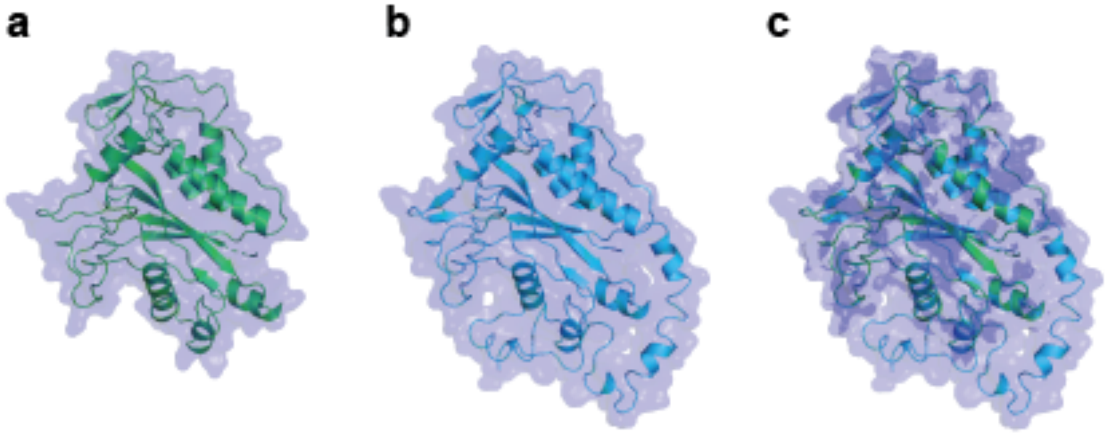


**Supplementary Fig. 11. a.** Protein structure of Cmr4 (PDB4W8W). **b.** Protein homology model of subtype IV-C Csf2 from *Thermococcus onnurineusfrom* (WP_012571280.1) generated with the Phyre2 protein structure prediction server (intensive mode); 81% (268 residues) of the query protein were modelled at >90% confidence. Proteins of known structure found to bear the highest homology to Csf2 are type III-A/B backbone subunits: Cmr4 from Pyrococcus furiosus (85% coverage, 22% identity and 99.5% confidence) and CRISPR type III-associated RAMP protein Csm3 (83% coverage, 31% identity and 97.1% confidence). **c.** Superimposed structures of Cmr4 and Csf2 generated by the PyMol molecular visualization software.

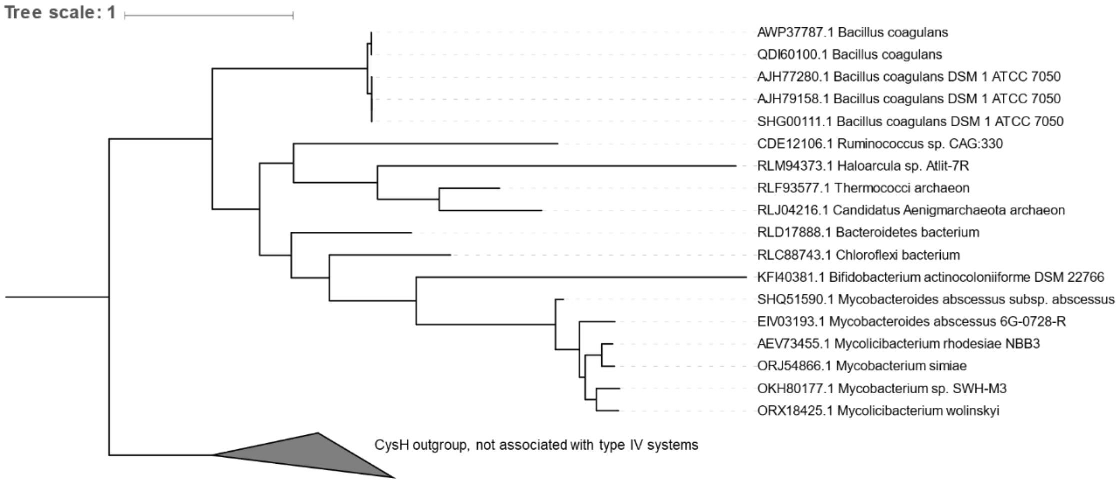


**Supplementary Fig. 12.** Phylogeny of CysH associated with subtype IV-B systems, built with a subset of representatives for which the protein accession number and host organism are given. Representatives of CysH proteins not associated with type IV CRIPSR-cas systems were used to root the maximum likelihood tree.

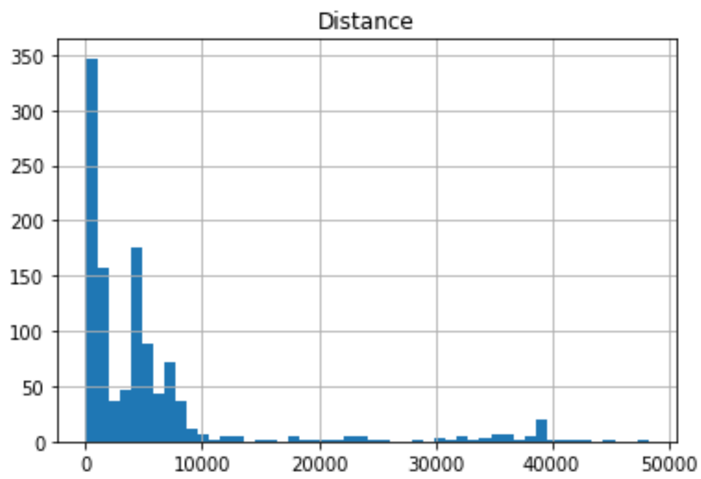


**Supplementary Fig. 13**. Distribution of distances between CRISPR arrays and the nearest cas operon. 10 kbp was used as a cutoff for pairing cas operons and CRISPR arrays.

### Supplementary Tables

**Supplementary Table 1.** HHpred-based protein homology and structure searches using the Cas10-like large subunit present in subtype IV as a query. It revealed significant matches to HD and zinc ribbon domains.

| | | | | | |
|---|---|---|---|---|---|
| **1** | PF01966 | 0.0046 | [ ------ ] | HD | HD domain. Metal dependent. |
| **2** | PF14353 | 0.022 | [ ----- ] | CpXC | CpXC protein. |
| **3** | PF09723 | 0.041 | [ ---- ] | Zn-ribbon_8 | Zinc ribbon domain. |
| **4** | PF05129 | 0.029 | [ ------ ] | Elf1 | Transcription elongation factor Elf1 like. |
| **5** | PF09538 | 0.075 | [ ----- ] | FYDLN_acid | Protein of unknown function (FYDLN_acid). |
| **6** | cd00077 | 0.058 | [ - ] | HDc | Metal dependent phosphohydrolase. |
| **7** | PF07514 | 00.23 | [ -- ] | TraI_2 | Putative helicase. |
| **8** | PF09706 | 00.25 | [ - ] | Cas_CXXC_CXXC | CRISPR-associated protein (Cas_CXXC_CXXC). |
| **9** | TF00277 | 00.31 | [ ------- ] | HDIG | HDIG domain. |
| **10** | PF05605 | 00.26 | [ ---- ] | zf-Di19 | Drought induced 19 protein (Di19), zinc-binding. |

**Supplementary Table 2.** HHpred-based protein homology and structure search results using the Csf1-like protein from type IV-A3 as a query.

| | | | | | |
|---|---|---|---|---|---|
| **1** | cd16341 | 0.0067 | [ -------- ] | FdhE | Formate dehydrogenase accessory protein FdhE. |
| **2** | PF13453 | 00.15 | [ --------- ] | zf-TFIIB | Transcription factor zinc-finger. |
| **3** | PF03264 | 00.12 | [ --------- ] | Cytochrom_NNT | NapC/NirT cytochrome c family, N-term. |
| **4** | PF11672 | 00.35 | [ ------- ] | DUF3268 | zinc-finger-containing domain. |
| **5** | PF09723 | 0.044 | [ ------- ] | Zn-ribbon_8 | Zinc ribbon domain. |
| **6** | cd14089 | 01.02 | [ -------- ] | STKc_MAPKAPK | Catalytic domain of the Serine/Threonine. |
| **7** | PF14446 | 02.05 | [ -------- ] | Prok-RING_1 | Prokaryotic RING finger family 1. |
| **8** | PF11845 | 01.02 | [ -------- ] | DUF3365 | Protein of unknown function (DUF3365). |
| **9** | PF06689 | 01.02 | [ ------- ] | zf-C4_ClpX | ClpX C4-type zinc finger. The ClpX heat shock. |
| **10** | cd06266 | 20 | [ -------------------- ] | RNase_HII | Ribonuclease H (RNase H) type II family. |

**Table 3.** HHpred-based protein homology and structure search results using the Csf3-Csf1 fusion protein from subtype IV-E as a query.

| | | | | | |
|---|---|---|---|---|---|
| **1** | TF03116 | 4.7E-25 | [ -------- ] | cas5_csf3 | CRISPR type IV/AFERR-associated protein |
| **2** | cd09707 | 4.8E-25 | [ -------- ] | Csf3_U | CRISPR/Cas system-associated RAMP superfamily |
| **3** | cd09705 | 0.063 | [ ------- ] | Csf1_U | CRISPR/Cas system-associated protein Csf1. |
| **4** | TF03114 | 0.063 | [ ------- ] | cas8u_csf1 | CRISPR type AFERR-associated protein Csf1 |
| **5** | PF10040 | 00.11 | [ -- ] | DUF2276 | Uncharacterized conserved protein (DUF2276). |
| **6** | PF03787 | 00.04 | [ --- ] | RAMPs | RAMP superfamily. |
| **7** | cd09652 | 01.01 | [ --- ] | Cas6-I-III | CRISPR/Cas system-associated RAMP superfamily. |
| **8** | TF01877 | 01.06 | [ --- ] | cas_cas6 | CRISPR-associated endoribonuclease Cas6. |
| **9** | cd16656 | 02.09 | [ -- ] | RING-Ubox_PRP19 | U-box domain, a modified RING finger. |
| **10** | TF01898 | 08.02 | [ ---- ] | cas_TM1791_cmr6 | CRISPR type III-B/RAMP module RAMP. |

**Supplementary Table 4.** Spacer-protospacer match analysis performed for all detected CRISPR-Cas subtypes and variants.

| | Plasmids | Viruses | Both | Unknown | Total |
|---|---|---|---|---|---|
| **All** | 225 | 336 | 60 | 7208 | 7829 |
| **Type IV** | 98 | 16 | 9 | 928 | 1051 |
| **Others** | 127 | 320 | 51 | 6280 | 6778 |
| **I-B** | 12 | 31 | 3 | 1607 | 1653 |
| **I-C** | 7 | 10 | 0 | 889 | 906 |
| **I-D** | 0 | 0 | 0 | 133 | 133 |
| **I-E** | 69 | 104 | 38 | 1756 | 1967 |
| **I-F** | 32 | 163 | 10 | 914 | 1119 |
| **I-U** | 0 | 1 | 0 | 72 | 73 |
| **II-C** | 1 | 1 | 0 | 36 | 38 |
| **III-A** | 1 | 4 | 0 | 544 | 549 |
| **III-B** | 4 | 0 | 0 | 152 | 156 |
| **III-D** | 1 | 6 | 0 | 177 | 184 |
| **IV-A1** | 9 | 2 | 0 | 350 | 361 |
| **IV-A2** | 6 | 0 | 0 | 174 | 180 |
| **IV-A3** | 79 | 13 | 9 | 200 | 301 |
| **IV-B** | 0 | 0 | 0 | 8 | 8 |
| **IV-C** | 0 | 0 | 0 | 26 | 26 |
| **IV-D** | 0 | 1 | 0 | 48 | 49 |
| **IV-E** | 4 | 0 | 0 | 122 | 126 |
| **Total** | **225** | **336** | **60** | **7208** | **7829** |

#### Supplementary Data

**Supplementary Data 1** Rare *cas6* genes associated to subtype IV-B loci.

**Supplementary Data 2.** Predictions of the genomic context of the identified type IV loci: plasmid-like elements or (pro)phages/viruses predictions.

**Supplementary Data 3.** Type IV Spacers-protospacer match analysis results against PLSDB. List of all detected non-coding/unknown ORF/gene family hits.

**Supplementary Data 4.** Type IV Spacers-protospacer match analysis results against the virus database. List of all detected non-coding/unknown ORF/gene family hits.

**Supplementary Data 5.** Rare cases of adaptation module components (Cas1/Cas2) found adjacent to type IV systems.
